# A dataset of 5 million city trees: species clustering and climate effects in urban forests

**DOI:** 10.1101/2022.03.18.484862

**Authors:** Dakota E. McCoy, Benjamin Goulet-Scott, Weilin Meng, Bulent Furkan Atahan, Hana Kiros, Misako Nishino, John Kartesz

## Abstract

Sustainable cities depend on urban forests. City trees improve our health, clean the air, store CO_2_, and cool local temperatures. Comparatively less is known about urban forests as ecosystems, particularly their spatial composition, nativity statuses, biodiversity, and tree health. Here, we assembled and standardized a new dataset of N=5,132,890 trees from 63 of the largest US cities with detailed information on location, health, nativity status, and species. We further designed new tools to analyze the ecosystem structure of urban forests, including spatial clustering and abundance of native trees, and validate these tools in comparison to past methods. We show that city trees are significantly clustered by species in 93% of cities, potentially increasing pest vulnerability (even in cities with biodiverse urban forests). Further, non-native species significantly homogenize urban forests across cities, while native trees comprise 0.44%-85.6% (median=45.6%) of city tree populations. Native trees are less frequent in drier cities, and indeed climate significantly shapes both nativity and biodiversity in urban forests. Parks are more biodiverse than urban settings. Compared to past work which focused primarily on canopy cover and species richness, we show the importance of analyzing spatial composition and nativity statuses in urban forests (and we created new datasets and tools to do so). This dataset could be analyzed in combination with citizen-science datasets on bird, insect, or plant biodiversity; social and demographic data; or data on the physical environment. Urban forests offer a rare opportunity to intentionally design biodiverse, heterogenous, rich ecosystems.

## Introduction

Cities are ecosystems. Humans (Willis & Petrokofsky, 2017) other animals (Berthon et al., 2021) depend on urban forests for many services. City trees improve mental and physical health (Hartig & Kahn Jr, 2016), capture and store carbon dioxide (Rowntree & Nowak, 1991), scrub toxic particulate matter from the air (Nowak et al., 2014), and cool local temperatures by about 0.83°C for every 10% increase in forest cover (Kong et al., 2014). The financial benefits of having a tree-rich city-- rather than a concrete jungle-- are huge and well-documented (McPherson et al., 2016). Tree inventories provide a wealth of useful data (Cowett & Bassuk, 2014, 2020; Galle et al., 2021; Kendal et al., 2014; McPherson et al., 2016; Richards, 1983; Steenberg, 2018). Many studies underscore the importance of city trees to humans, but comparatively fewer evaluate urban forests as potentially biodiverse ecosystems (Alvey, 2006). Through this ecological lens, it is important to understand species diversity (Behm, 2020), nativity status (Tallamy, 2004), and spatial arrangements (Roman et al., 2018) of city trees.

Here, we assembled a dataset of N=5,132,890 individual trees from 63 US cities (Data S1) with precise data on species, exact location, native status, and standardized condition. We also developed tools to analyze the biodiversity, spatial structure, native tree abundance, and overall condition of urban forests. We demonstrate that these new tools provide a richer picture of urban forests than relying on canopy cover and species richness alone. For example, it is now possible for researchers to assess the spatial arrangement of trees by species (taking into consideration the underlying spatial arrangement of city streets)—a metric which, we show, is not dependent on biodiversity and which may indicate vulnerability to pests such as Dutch Elm disease (Laćan & McBride, 2008). Likewise, we show that the abundance of native trees varies greatly (even among cities with a high biodiversity of trees); native tree abundance is a useful proxy for an environment’s capacity to support diverse communities of birds, butterflies, and other animals (Burghardt et al., 2009, 2010; Tallamy, 2004). Taken together, we make available a large new dataset of city trees; user-friendly tools to better analyze the ecosystem structure of urban forests; and proof-of-concept analyses to demonstrate potential uses of the data. Through these technical and practical advances, we take a first step toward enabling the design of rich, heterogenous ecosystems built around urban forests.

## Results and Discussion

### A new dataset of more than 5 million city trees

First, we assembled and standardized a large dataset of N=5,132,890 city trees to enable the analysis of urban forests’ ecosystem structure. We acquired street tree inventories from 63 of the largest 150 US cities (those which had conducted inventories) and developed a standardization pipeline in *R* and *Python* (Data S9). Each inventory was produced using different, city-specific methods: for example, some cities only reported a tree’s common name; some reported an address but no coordinates; some reported tree size in feet, some in meters; some scored tree health from 1-5 while others rated trees as “good” or “poor;” etc. Very few cities reported a tree’s native status. Therefore, we inspected metadata for all cities (and communicated with urban officials) to standardize column names, standardize metrics of tree health, and convert all units to metric (Table S4). We converted all common names to scientific and manually corrected misspellings in all species names (see Data S9, and Materials and Methods, for full details). We manually coded all tree locations as being in a green space or in an urban environment to enable comparisons between location types. Finally, we referenced data from the Biota of North America Project on native status to classify each tree as native or not. The resulting dataset (Figure 1, Data S1) comprised 63 city datasheets each with 28 standardized columns (Table S4).

**Fig. 1.**
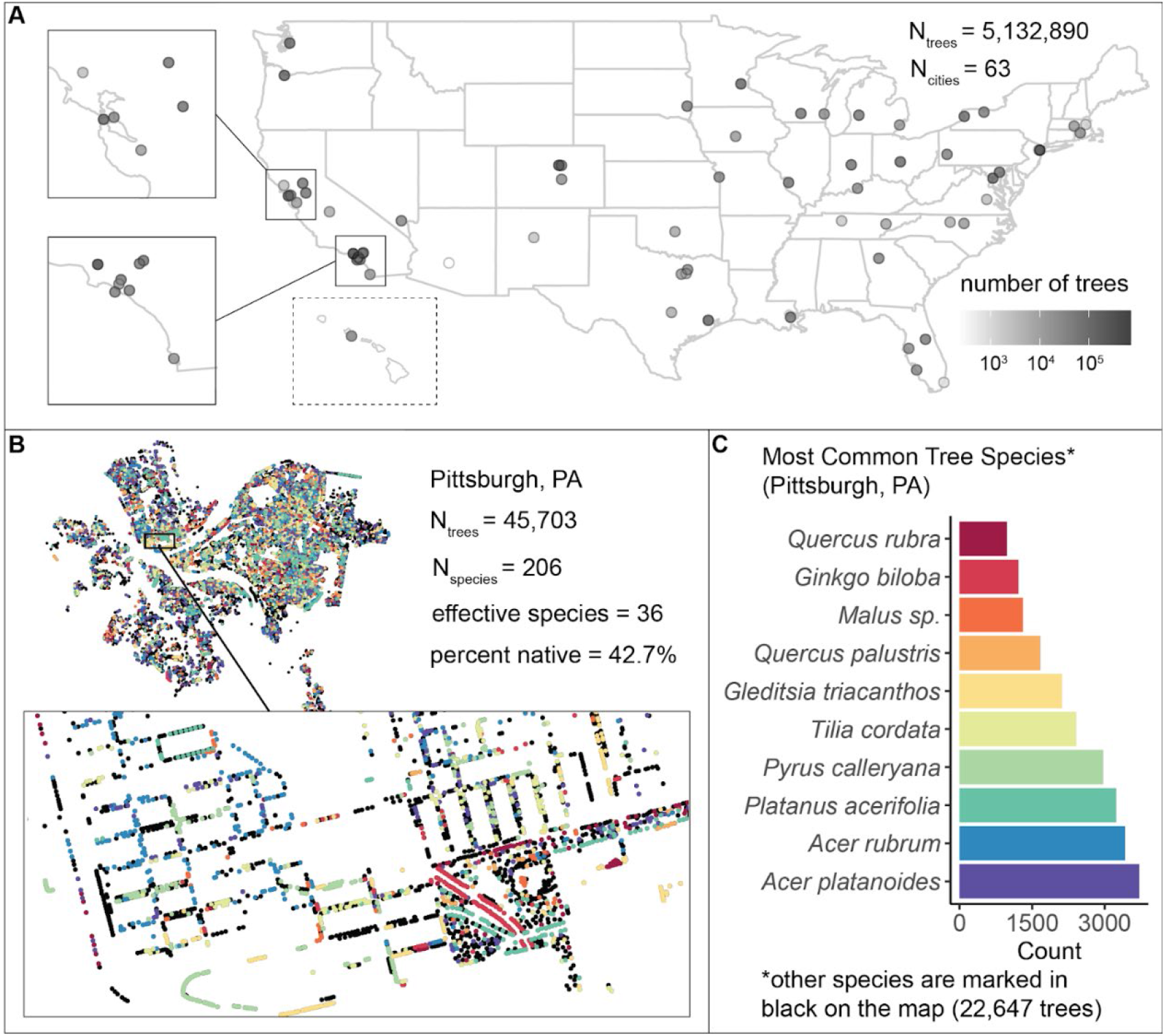
We assembled and standardized a dataset of N=5,132,890 street trees from publicly available street tree inventories across 63 cities in the USA. **(A)** The number of trees recorded per city varied from 214 (Phoenix, AZ) to 720,140 (Los Angeles, CA) with a median of 39,193. **(B)** Sample plot of Pittsburgh, PA with trees colored by species type (inset: zoomed-in view of trees lining streets and parks). **(C)** Counts of the ten most common species inventoried in Pittsburgh; not shown are 22,647 trees belonging to other species (black points in (B)). The dataset includes information on species, exact location, whether or not a tree is native, tree height, tree diameter, location type (green space or urban setting), tree health/condition, and more (Data S1). Source data is Data S1 and Data S2; source code is Data S4.

### New tools for-- and preliminary analyses of-- biodiversity, spatial structure, native status, tree health, and climate effects

Urban forests are typically quantified through species richness (as a measure of biodiversity) and percent canopy cover. Our large, fine-grained dataset allows for analysis of (i) effective species counts (a measure that allows comparison between cities of different sizes), (ii) spatial structure of city tree communities, (iii) abundance of native trees, (iv) climate drivers of biodiversity and native tree abundance, and (v) how urban forest biodiversity correlates with fine-grained data on socio-economics, demographics, the physical environment, and other forms of biodiversity such as birds and insects.

We found that city tree communities are moderately biodiverse, particularly in parks (Fig. 2), but are significantly clustered by individual species (Fig. 3). Urban forests varied in number of species represented (min=16, median=137, max=528; Data S2) and in a robust, naturalistic measure of biodiversity known as effective species count (min=6, median=26, max= 68; Fig. 2A). Trees located in parks were significantly more biodiverse than trees located in developed environments (e.g., along streets; Fig. 2B, Fig. S1).

**Fig. 2:**
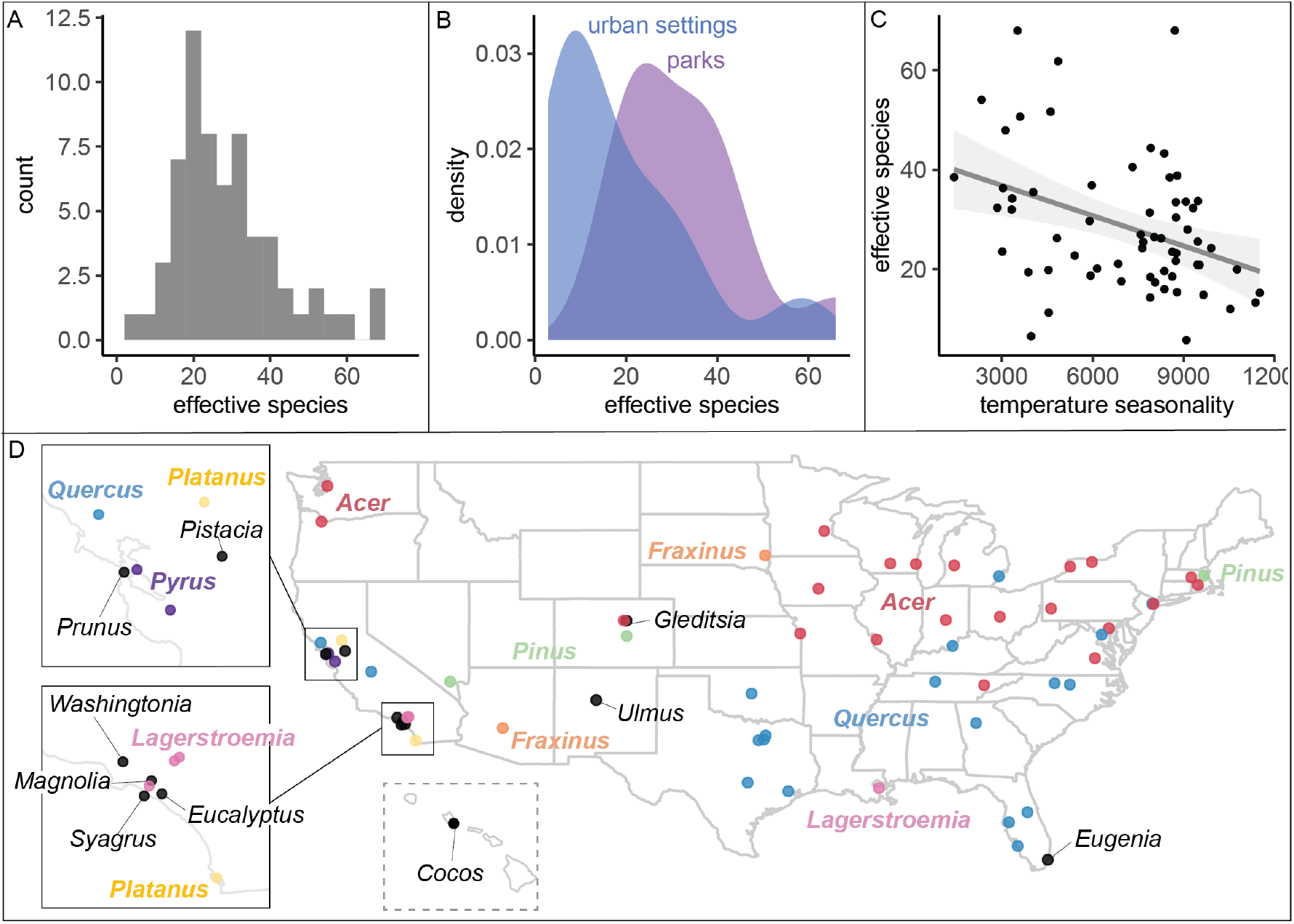
City tree communities are biodiverse and shaped by climate, although certain genera dominate. **(A)** Effective species count, a measure of biodiversity, ranged across cities from min=6 to max=58 with a median=26. Effective species count is a more nuanced metric than abundance-based metrics (supporting Fig. S2). **(B)** Trees in parks were significantly more biodiverse than trees in urban settings such as along streets (two-sample paired t-test; t=7.2, p<0.0005, 95% CI=[9.99, 18.6], mean diff. = 14.3; see supporting Fig. S1). **(C)** Environmental factors were significantly correlated with effective species count but socio-cultural variables were not (supporting Table S1); biodiversity is negatively related to temperature seasonality (captured through environmental PC1). **(D)** The most abundant genus in each city is labeled here; see most common species by city in supporting Fig. S3. Supporting figures for this figure include Fig. S1, Fig. S2, and Fig. S3; Table S1 is a supporting table. Source data is Data S1 and Data S2; source code is Data S4; and an associated tool to calculate effective species is Data S3.

**Fig. 3:**
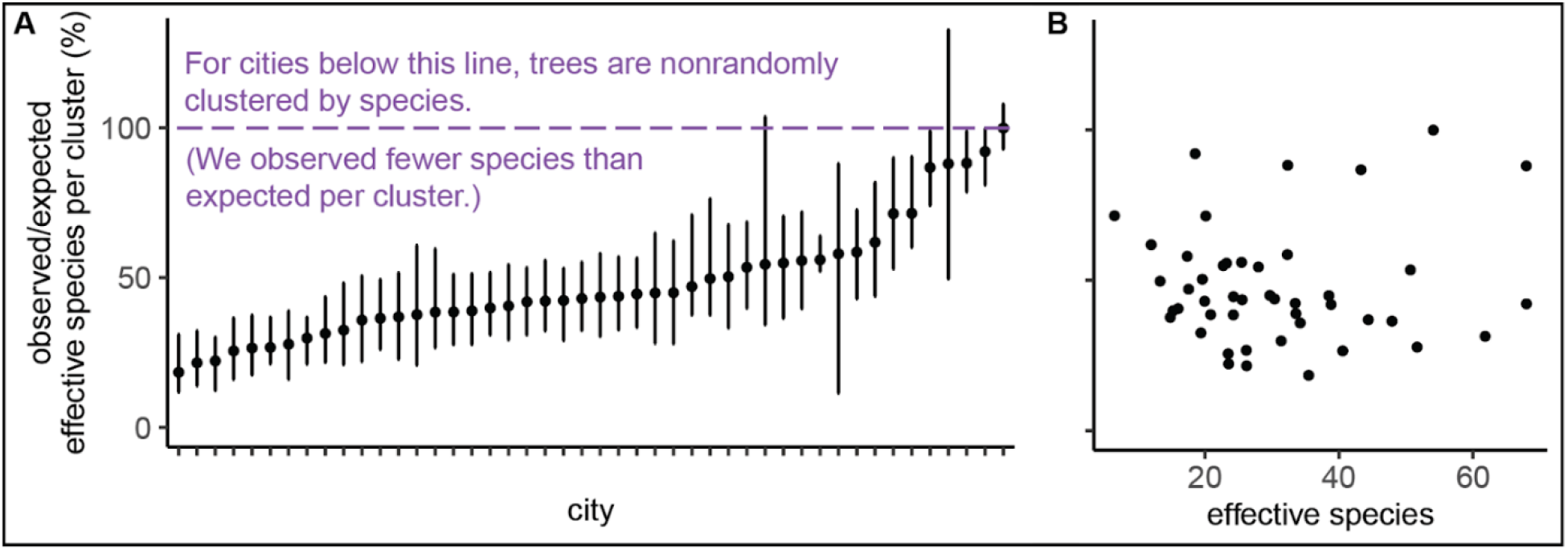
Trees are spatially clustered by species in nearly all cities, even in cities with high biodiversity. **(A)**. In 43 of 46 cities, trees are non-randomly clustered by individual species (with significantly fewer effective species per spatial cluster than expected, values<100%). Plotted points represent median values and IQRs (observed / expected effective species counts) for all clusters in a city (see N_clusters_ per city and full statistics in Data S5). **(B)** The degree of **s**patial clustering in a city was not correlated with biodiversity (supporting Fig. S4). N_cities_=46. Fig. S4 is a supporting figure for this figure. Source data is Data S1 and Data S5; source code is Data S4.

Another commonly used biodiversity metric is maximum abundance. Many foresters follow Santamour’s 10/20/30 rule: plant no more than 10% trees of the same species, 20% genus, and 30% family (Santamour Jr, 2004). Maximum abundance per species correlated significantly with effective species but cities below the 10% max abundance threshold vary from ~20-70 effective species (Fig. S2). Therefore, Santamour’s rule may be a necessary but not a sufficient guideline for urban forests, so we developed an Excel resource to calculate effective species from a list of (i) species counts or (ii) all trees (Data S3).

Because our dataset spans many different environmental conditions, we could assess the extent to which climate has impacted the ecosystem structure of city trees. Across the USA, climate--but not socio-cultural factors-- correlated with city tree biodiversity (Fig. 2C, Table S1). Specifically, temperature and rainfall significantly correlate with effective species count (aligning with previous analyses of city trees (*17*) and global distributions of plants (*21*)). Maples (*Acer*) and Oaks (*Quercus*) dominated city tree genera across the country (Fig. 2D), while the most common species were *Acer platanoides* (Norway Maple), *Fraxinus pennsylvanica* (Green Ash), *Lagerstroemia indica* (crape myrtle), and *Platanus acerifolia* (Fig. S3).

We next investigated the spatial arrangement of biodiversity in urban forests. Biodiverse, rather than species-poor, city tree communities offer many well-documented benefits. Biodiverse forests are more effective in resisting diseases (Laćan & McBride, 2008), are more resilient in the face of climate change (Roloff et al., 2009) and confer greater mental health benefits (Fuller et al., 2007). Compared to biodiversity, the spatial arrangement of trees is less well-understood, even though clusters of same-species of trees may be more susceptible to pest outbreaks (Greene & Millward, 2016; Raupp et al., 2006).

We found that city trees were nonrandomly clustered by individual species in 43 of 46 cities (Fig. 3A). Additionally, a city’s clustering score was not significantly correlated with biodiversity metrics and is therefore a separate metric of interest (Fig. 3B, Fig. S4). Urban forests with well-mixed arrangements of trees may be more resistant to species-specific diseases and blights, as in the case of the Emerald Ash Borer *Agrilus planipennis* (Greene & Millward, 2016). Clustering by species is not necessarily a negative; further research could consider the causes and consequences of clustering by species in urban forests. As city officials consider which trees to plant where, weighing many factors such as aesthetics and hardiness (Conway & Vander Vecht, 2015), we suggest they consider a simple metric of species clustering. To calculate clustering metrics, readers familiar with Python and R can use the code in Data S4; others should contact the authors (a web resource is currently under development).

Our new dataset allows researchers and urban foresters to consider the utility of native vs. non-native trees. Whether or not a city decides to plant native as opposed to introduced species is a growing topic of interest (along with whether nativity matters, and how to define “nativity” (Berthon et al., 2021; Gould, 1998; Sjöman et al., 2016)). We classify plants as “native” if they occur in a particular region without direct or indirect human intervention. This definition does not account for the substantial effects of Indigenous peoples on plant communities before European contact, nor does this paper address the flaws with a “native-or-not” ecological approach (see discussion of an alternative Indigenous ecology in (Grenz, 2020; McKay, 2021)).

Here we found that the percent of trees that were native varied across cities from 0.44% to 85.6% with a median of 45.6% (Fig. 4). Wetter, cooler climates correlated with significantly higher percentages of native trees (Fig. 4A-B). However, it is important to note a strong east-to-west gradient, by which fewer native trees were present in western states (Figure 4A). Thus, some social factor may have influenced the planting of native trees (Roman et al., 2018; Steenberg, 2018). However, after accounting for climate, younger cities had a higher percentage of native trees (Table S2); perhaps urban forestry practitioners have been more likely to consider nativity status in recent years. The observed east-to-west gradient deserves further research attention.

**Fig. 4:**
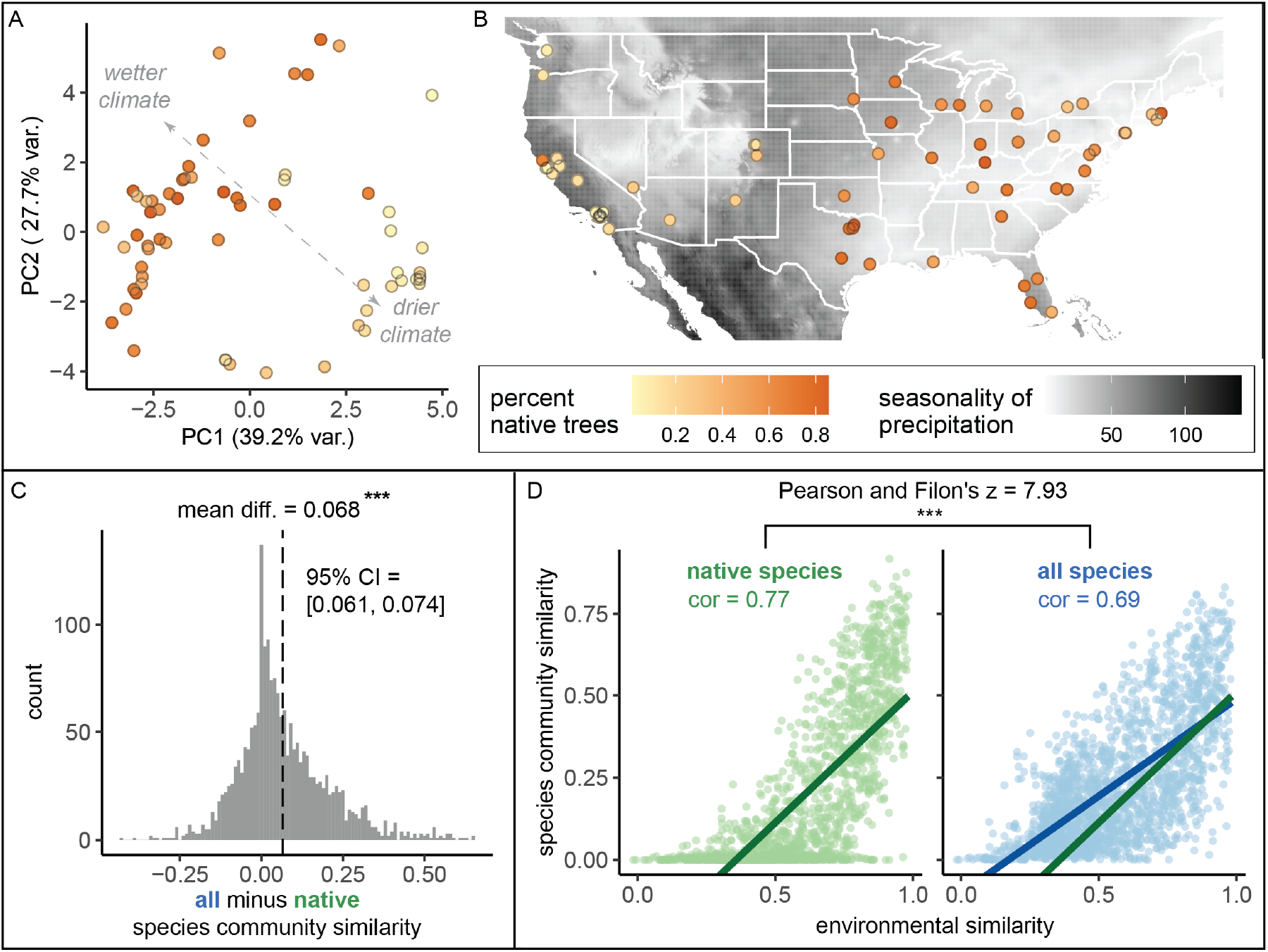
Environment strongly influences native trees, while non-native trees make species compositions more similar between cities. **(A)** Cities in wetter, cooler climates -- and younger cities -- had significantly higher percentages of native trees (beta regression; AIC= −61.3, pseudo-R^2^=0.64, log likelihood=36.7; statistics in Table S2). Here we plot a principal component analysis of the Bioclim variables (Data S8), colored by percent native trees. **(B)** The percent of native trees is plotted against seasonality of precipitation (black and white background). **(C)** Among city pairs (N_comparisons_=1953), overall species communities are significantly more similar to one another than their native species communities alone (paired t-test, t= 21.1, p < 0.0005; result upheld by non-parametric Wilcoxon signed rank test). We calculated chi-square similarity scores for each city pair for (i) native-only species and (ii) all species and subtracted (i) from (ii). **(D)** Among city pairs, environment is significantly more strongly related to native species than non-native. We compared chi-square similarity scores between species communities (left: native-only; right: all) against environmental similarity scores (one minus the normalized euclidean distance in our PCA). Left panel, green, native species only: Pearson’s product-moment correlation, cor = 0.77, 95% CI= [0.75, 0.78], t = 52.6, p<0.0005. Right panel, blue, all species: Pearson’s product-moment correlation; cor = 0.69, 95% CI= [0.66, 0.71], t = 42.0, p<0.0005). Fig. S5 is a supporting figure for this figure. Source data is Data S1, Data S2, Data S7, and Data S8; source code is Data S4; and an associated tool to check the native status of a list of tree species is Data S6.

In general, native plant species support richer local ecosystems (e.g., more diverse and numerous bird and butterfly communities (Burghardt et al., 2009, 2010). Among non-native plants, those with native congeners support more and more diverse Lepidopteran species than those without (Burghardt et al., 2010). Many cities with relatively low populations of native trees nonetheless had many non-native congeners of native species (bottom right quadrant, Fig S5B)—and therefore likely provide moderate insect habitat. Native tree biodiversity is significantly correlated with overall biodiversity (Fig. S5). Nativity status is a useful proxy for ecological value (although it is not, alone, a deciding factor (Berthon et al., 2021)), so we developed an Excel tool to report nativity status based on a submitted list of species for a given city or state (Data S6). Original BONAP data on all native taxa for each US state is available in Data S7.

Urban foresters must consider tree hardiness when selecting species to plant, and our dataset provides standardized metrics of tree health across many cities. Our preliminary analyses suggest that whether or not a tree was native had no clear impact on tree health (Table S3). Trees are generally healthier (i.e., have better condition) when they are smaller and/or in an urban setting rather than in parks (Table S3), possibly because city arborists quickly remove unhealthy trees in densely populated areas where they pose a fall risk. Further work is needed on within-species trends.

Are city tree communities more similar to each other than we would expect based on geography and climate? Indeed, we found that non-native trees drive similar species compositions between cities (Fig. 4C), reflecting the phenomenon of “biotic homogenization” (M. L. McKinney & Lockwood, 1999). Unsurprisingly, environment is a significant driver of tree community similarity between cities, but this association is stronger for native trees (Fig. 4D).

### Potential future analyses: socio-economics, demographics, the physical environment, and citizen-science biodiversity identification

Beyond the analyses demonstrated above, our dataset could also be combined with social, economic, and physical variables for new analyses (Fig. 5). Simple maps of biodiversity in the Washington, DC area (Fig. 5A-B) show that high biodiversity qualitatively overlaps with high median household income (Fig. 5C). In other words, not only do “trees grow on money”(Schwarz et al., 2015) but they may be more biodiverse in richer areas (Pedlowski et al., 2002). Biodiverse green spaces improve mental health more than species-poor spaces (Wood et al., 2018) and likely have other synergistic benefits (e.g., promoting more biodiversity among birds and insects); therefore, further analyses of urban forest biodiversity by income, and other demographic factors, would be useful.

**Fig. 5:**
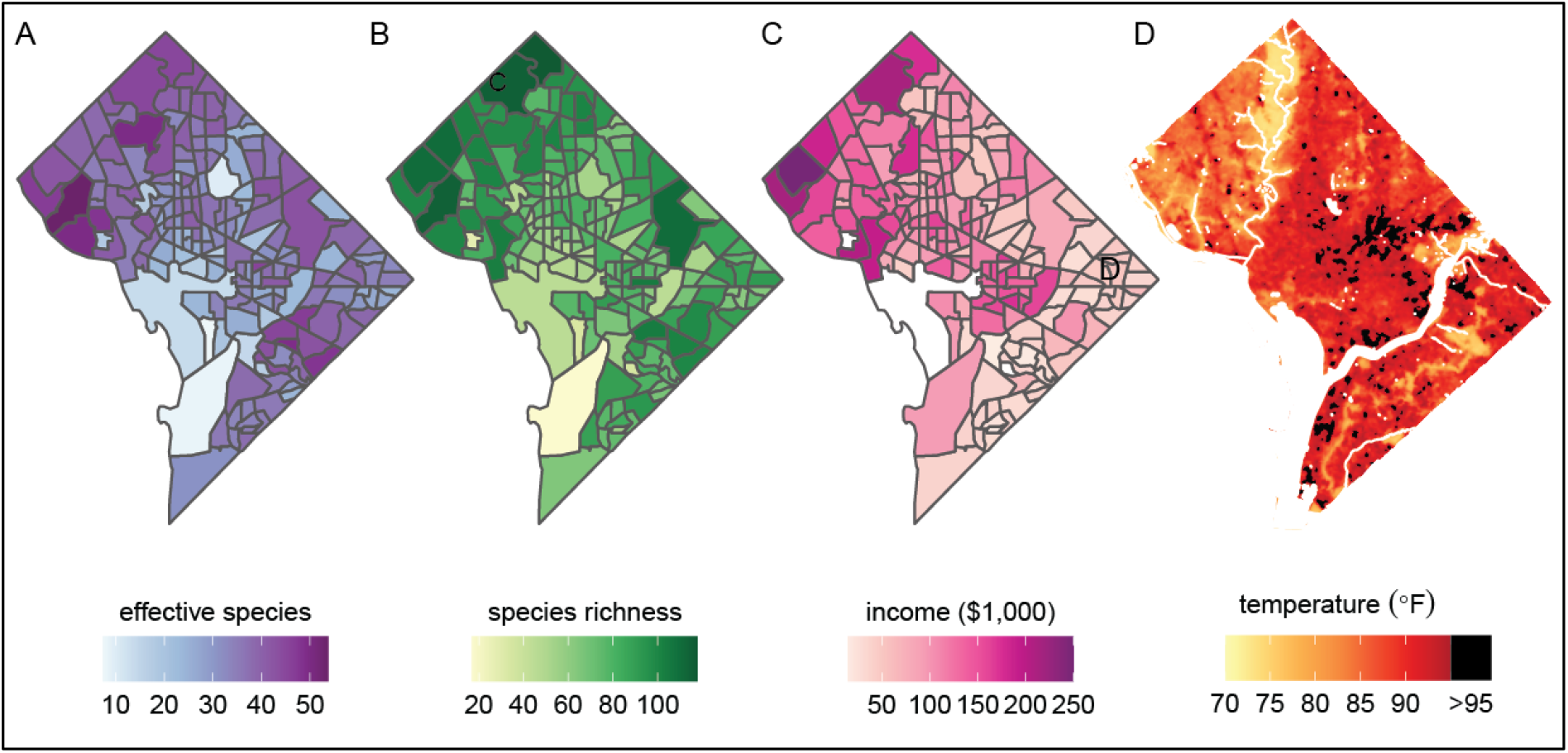
Future analyses could combine this city trees data with social, demographic, or physical variables (including income and urban heat islands). Here we plot different variables for Washington, DC, showing qualitative concordance between (A-B) measures of biodiversity, (C) household income, and (D) the location of urban heat islands. **(A)** Effective species count is highest in the northwest and varies by census tract from 7 species to 54 species (median = 35 species). **(B)**. Species richness is also highest in the northwest and varies by census tract from 17 species to 118 species (median 77 species). **(C)** Median household income is highest in the northwest, the region which overlaps substantially with the most biodiverse urban forests. **(D)** Land surface temperature in July 2018 shows that the highest temperatures, and urban heat islands with temperatures > 95°F, tend to overlap with less-richly-forested areas. Source data is Data S1 and open-access data available from the US Census and the DC Open Data Portal (see Materials and Methods) and source code is Data S4.

Urban forests cool city temperatures (Kong et al., 2014), another benefit which is not equitably distributed; indeed, Fig. 5D shows that heat islands are more common in species-poor (and less well-forested) areas of Washington, DC. The dataset herein could be combined with many physical variables for new analyses of temperature, air quality, and more.

This city trees dataset could also be analyzed in combination with other biodiversity datasets gathered by citizen-scientists. Members of the public frequently use popular phone applications to identify and document the location of birds, plants, insects, and more (Bonnet et al., 2020; Chandler et al., 2017). Future work could analyze whether a biodiverse urban forest correlates with a more biodiverse community of insects, birds, and even non-tree plants. Likewise, an analysis could consider whether the abundance of native trees correlates with other important measures of ecosystem health (such as insect abundance). Since citizen-science datasets typically include exact location, future work could assess these trends over fine scales (e.g., within particular parks or in bounded neighborhoods) as well as across cities.

### Conclusion

Urban forests are ecosystems over which humans exercise precise control. We have an opportunity to design city tree communities that are biodiverse, spatially heterogenous, and include native species—thereby building resilience against climate change (Roloff et al. 2009), avoiding pest/pathogen outbreaks (Laćan & McBride, 2008), improving human’s mental and physical health (Fuller et al., 2007), and providing richer habitat for non-human animals (Burghardt et al., 2009, 2010; Gallo & Fidino, 2018; Parsons et al., 2018). We should use biodiversity-informed decision-making to forge a path toward a green urban future.

## Materials and Methods

### Data Acquisition

We limited our search to the 150 largest cities in the USA (by census population). To acquire raw data on street tree communities, we used a search protocol on both Google and Google Datasets Search (https://datasetsearch.research.google.com/). We first searched the **city name** plus each of the following: **street trees, city trees, tree inventory, urban forest**, and **urban canopy** (all combinations totaled 20 searches per city, 10 each in Google and Google Datasets Search). We then read the first page of google results and the top 20 results from Google Datasets Search. If the same named city in the wrong state appeared in the results, we redid the 20 searches adding the state name. If no data were found, we contacted a relevant state official via email or phone with an inquiry about their street tree inventory. Datasheets were received and transformed to .csv format (if they were not already in that format). We received data on street trees from 64 cities. One city, El Paso, had data only in summary format and was therefore excluded from analyses.

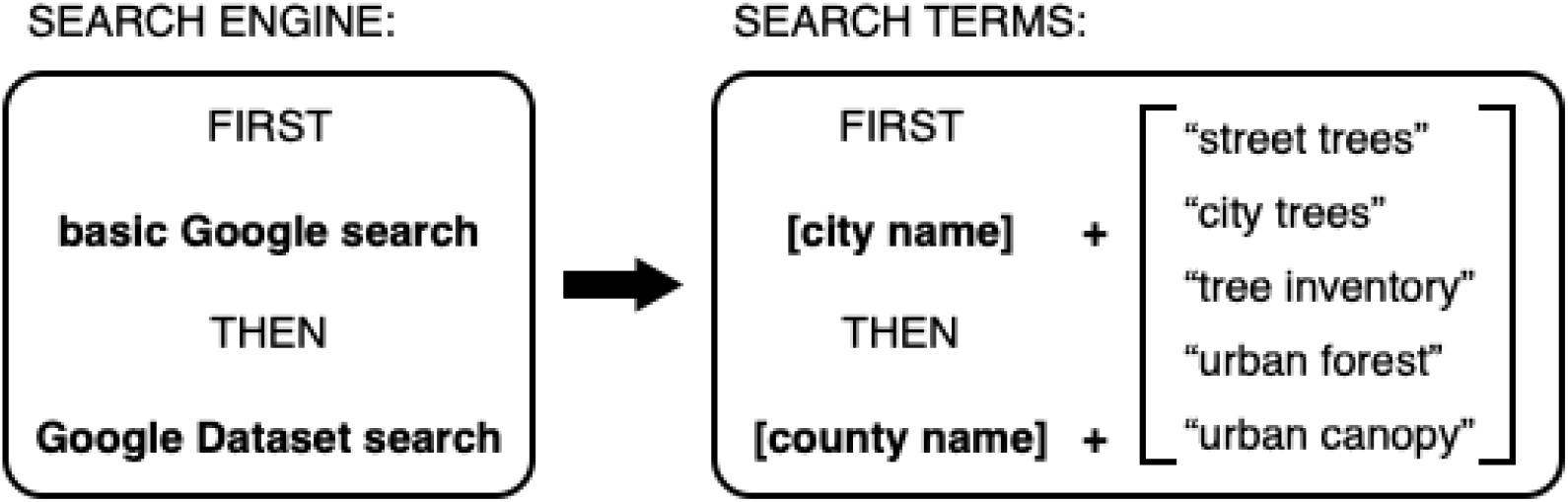

### Data Cleaning

All code used is in the zipped folder Data S5. Before cleaning the data, we ensured that all reported trees for each city were located within the greater metropolitan area of the city (for certain inventories, many suburbs were reported - some within the greater metropolitan area, others not).

First, we renamed all columns in the received .csv sheets, referring to the metadata and according to our standardized definitions (Table S4).

Second, we used pandas in Python (W. McKinney & Others, 2011) to correct typos, non-ASCII characters, variable spellings, date format, units used (we converted all units to metric), address issues, and common name format. In some cases, units were not specified for tree diameter at breast height (DBH) and tree height; we determined the units based on typical sizes for trees of a particular species. Wherever diameter was reported, we assumed it was DBH. We created a column called “location_type” to label whether a given tree was growing in the built environment or in green space. All of the changes we made, and decision points, are preserved in Data S9.

Third, we checked the scientific names reported using gnr_resolve in the R library taxize (Chamberlain & Szöcs, 2013), with the option Best_match_only set to TRUE (Data S9). Through an iterative process, we manually checked the results and corrected typos in the scientific names until all names were either a perfect match (n=1771 species) or partial match with threshold greater than 0.75 (n=453 species). BGS manually reviewed all partial matches to ensure that they were the correct species name, and then we programmatically corrected these partial matches (for example, *Magnolia grandifolia*-- which is not a species name of a known tree-- was corrected to *Magnolia grandiflora*, and *Pheonix canariensus* was corrected to its proper spelling of *Phoenix canariensis*). Because many of these tree inventories were crowd-sourced or generated in part through citizen science, such typos and misspellings are to be expected.

Some tree inventories reported species by common names only. Therefore, our fourth step in data cleaning was to convert common names to scientific names. We generated a lookup table by summarizing all pairings of common and scientific names in the inventories for which both were reported. We manually reviewed the common to scientific name pairings, confirming that all were correct. Then we programmatically assigned scientific names to all common names (Data S9).

Fifth, we assigned native status to each tree through reference to the Biota of North America Project (Kartesz, 2018), which has collected data on all native and non-native species occurrences throughout the US states. Specifically, we determined whether each tree species in a given city was native to that state, not native to that state, or that we did not have enough information to determine nativity (for cases where only the genus was known).

Sixth, some cities reported only the street address but not latitude and longitude. For these cities, we used the OpenCageGeocoder (https://opencagedata.com/) to convert addresses to latitude and longitude coordinates (Data S9).

Seventh, we trimmed each city dataset to include only the standardized columns we identified in Table S4.

After each stage of data cleaning, we performed manual spot checking to identify any issues.

### Environmental Variables

We retrieved WorldClim data on 19 climate variables using the *getData* function in package *raster* (Hijmans & van Etten, 2012) with parameters var="bio” and res=2.5. We gathered climate variables for each city by extracted the grid cell closest to the latitude and longitude of each city in our dataset, and then performed a PCA on the environmental variables.

### Biodiversity

We calculated effective species counts (the exponent of the Shannon-Weiner index) as our measure of biodiversity because it incorporates both richness (number of species) and evenness (distribution of those species; (Kendal et al., 2014), and because it is a metric that behaves naturally and intuitively in comparisons between species communities (see http://www.loujost.com/Statistics%20and%20Physics/Diversity%20and%20Similarity/EffectiveNumberOfSpecies.htm). Effective species count is calculated as shown in Equation 1, where *n* is the number of species present and *p_i_* is the frequency of a species *i*.

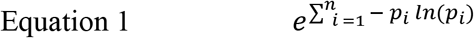

To determine what environmental and socio-cultural factors drive biodiversity (dependent variable: effective species count), we used the *olsrr* package in R (Hebbali & Hebbali, 2017) to compare AIC and adjusted R2 values for all possible models incorporating the following independent variables: environmental PCA1, environmental PCA2, environmental PCA1 * environmental PCA2, city age, tree city USA (whether or not a city was designated as a tree city USA), city age * tree city USA, and the log-transformed number of trees in a given city. We identified the best-fitting four models and report statistics in Table S1. We ran all models with and without the two strongest outlier cities, Miami, FL and Honolulu, HI.

### Spatial Structure

We wanted to quantify the degree to which trees were spatially clustered by species within a city (rather than randomly arranged). To do so, we first clustered all trees within each city using hierarchical density based spatial clustering through the *hdbscan* library in Python (McInnes et al., 2017). *HDBSCAN*, unlike typical methods such as “k nearest neighbors”, takes into account the underlying spatial structure of the dataset and allows the user to modify parameters in order to find biologically meaningful clusters. For city trees, which are often organized along grids or the underlying street layout of a city, this method can more meaningfully cluster trees than merely calculating the meters between trees and identifying nearest neighbors (which may be close as the crow flies but separated from each other by tall buildings).

We converted latitude and longitude values within a city to their planar projection equivalents (in Universal Transverse Mercator (UTM)) using the from_latlon function in Python package UTM (Bieniek et al., 2016). In total we had N = 59 cities with spatial information about their trees.

We then clustered all the trees in a given city using *HDBSCAN* with parameters min_cluster_size=30, min_samples=10, metric=’manhattan’, cluster_selection_epsilon=0.0004, cluster_selection_method = ’eom’); we arrived at these parameters through trial and error with a sample set of cities.

Once we had all trees in a city assigned to spatial clusters (or, for trees far from the clusters, notated as “noise” and eliminated from further analysis), we used a bootstrapping method to quantify the degree of homogenization within spatial clusters. For each cluster of trees (e.g., a cluster of 120 trees in Pittsburgh, PA) we (i) calculated the observed effective species number; (ii) we randomly resampled 120 trees from Pittsburgh’s entire 45,703-tree-dataset and calculated the effective species number of that random group of 120 trees; (iii) we repeated step (ii) 500 times; (iv) we recorded the mean, median, and interquartile range of effective species counts from those 500 samples; and (v) we divided the expected effective species (median effective species count from all 500 samples) by the observed effective species count in the actual spatial cluster of 120 trees. The resulting value therefore quantifies the degree to which a spatial cluster is a random set of that city’s tree species (values close to 100%) or a nonrandom set of same-species clusters (values less than 100%).

### Nativity Status

To determine whether or not a tree was native to the state in which it appeared, we referred to the state-specific lists of native species from the Biota of North America Project. Each tree species was therefore coded as native = TRUE, native = FALSE, or native = no_info. Some tree records included only genus-level data, which was coded as “no_info”.

We performed beta regression models with a logit link function using the package betareg in R (Zeileis et al., 2019), with percent native trees in a given city as the dependent variable. We assumed the precision parameter ϕ did not depend on any regressors. We started with a model incorporating only environmental variables, based on the substantial evidence that climate impacts native species biodiversity, and then added one variable at a time to determine whether the additional variables improved the model’s performance (tested through the *lrtest()* function from the package *lmtest* (Hothorn et al., 2015). The best model incorporated the following dependent variables: environmental PCA1, environmental PCA2, log(number trees), and city age with no interaction terms. We identified one major outlier, Honolulu, HI, and re-ran the model excluding the outlier (results did not significantly change).

### Condition and Health

We asked whether a tree’s condition within a given city was correlated with size (DBH), location type (whether in the built environment or in green space such as a park), and nativity status. Fifteen cities had two or more of these variables with adequate sample sizes, and we ran separate logistic regression models by city because cities do not always score condition on comparable scales. We coded tree condition as a binary variable, where “excellent,” “good”, or “fair” condition trees were coded as 1 and “poor”, “dead”, and “dead/dying” trees were coded as 0. We used function *glm2()* in the R package *glm2* (Marschner c et al., 2011), and for each model determined whether it was a better fit than an empty model. We calculated odds ratios, confidence intervals, and p-values (see Table S3).

### Similarity Between Tree Communities

For N = 1953 city-city comparisons of street tree communities, we could calculate weighted measures of similarity because we had frequency data. We used chi-square distance metrics on species frequency data (because the actual count of trees reflected differences in sampling efforts between cities). Chi-square similarity is calculated following Equation 2, where n is the total number of species present in either city, *x* and *y* are vectors of species frequencies for the two cities being compared, and for each species *i*, *x_i_* is the frequency of that species in city *x* and *y_i_* the frequency of the same species in city *y*. Chi-square similarity is one minus the chi-square distance.

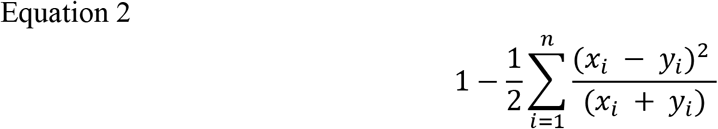

We calculated environmental similarity as one minus the normalized euclidean distance in our PCA plot of environmental variables.

To determine whether city species similarity was driven by native species, non-native species, or neither, we performed a two-sample paired t.test using the function *t.test* in R between the native species chi-squared similarity scores and the all-species chi-squared similarity scores. Because the variables were not perfectly normally distributed (although they were even and symmetric), we also performed a non-parametric Wilcoxon signed rank test. We plotted a histogram of the difference between each pair of city’s chi squared scores for (i) all species and (ii) native species only.

To determine whether the environment was a stronger driver of native species communities versus all species communities, we compared correlation scores. Specifically, we used the function *cor.test* in R to calculate the Pearson’s product-moment correlation between chi-squared similarity and environmental similarity for (i) native species only and (ii) all species. We compared all-species-environment correlation to the native-species-environment by calculating Pearson and Filon’s z using the *cocor* package in R (Diedenhofen & Musch, 2015) for two overlapping correlations based on dependent groups (calculation takes into account correlation between chisq_native and chisq_all, among other things).

### Income and Urban Heat Islands

To demonstrate the value of our dataset for analyses of social, economic, and physical variables, we mapped several such variables for Washington, DC using packages *raster* (Hijmans & van Etten, 2012), *sf* (Pebesma, 2018), and *tidycensus* (Walker et al., 2021) in R. First, we split our trees data by census tract and mapped species richness and effective species count within each tract; next, we extracted median household income data and plotted it for each census tract (Walker, 2022). Finally, we downloaded LANDSAT data on surface temperatures in DC for July 2018 from the DC Open Data portal (https://opendata.dc.gov/documents/land-surface-temperature-july-2018/explore; CC-BY-4.0) and plotted this, marking heat islands (temperature > 95°F) in black (Jolly, 2019).

## Acknowledgments

We are so grateful to the innumerable citizen-foresters, trained arborists, and city government officials who have worked tirelessly to make this data available. For their help with this specific project, we wish to thank Shane D. McQuillan (Des Moines, IA), Jane Gregory and Maria Repass (Orange County, FL), Bailey Patterson (Overland Park), Joran Viers (Albuquerque, NM), Brian Liberti (Rochester City, NY), Gretchen Erickson (Huntington Beach, CA), Randy Menzel (Huntington Beach, CA), Andrew Pineda (Huntington Beach, CA), Terri Bladow (Santa Rosa, CA), Matt Stull (Santa Rosa, CA), John Saylor (Lexington, KY), Todd Hayes (Greensboro, NC), Gary Farris (Wichita, KS), Donna Davis (Eastern Colorado), Dan Buckler (Wisconsin), Nathan Randolph (Baltimore, MD), Erik Dihle (Baltimore, MD), Daniel Hickey (Durham, NC), Glenn Slaton (Durham, NC), The City of Saint Louis Forestry Division, Steven Ashley (Louisville, KY), Kasey Krause (Knoxville, TN), Russell Calhoun Jr.(Houston, TX), David Wrights (Oklahoma City, OK), Donna Davis (CO), Rachot Moragraan (Garden Grove City, CA), Kevin Wilde and Bill Williams (Amarillo, TX). For the data for Hawaii, we wish to thank Heather McMillen, Wai Lee, and Terri-Ann Koike, and we wish to state that data used to help generate this report has been collected by the Citizen Forester Program; a collaborative project of the State and Private Forestry branch of the USDA Forest Service, Department of Agriculture, Region 5; the Kaulunani Urban and Community Forestry Program of the DLNR Division of Forestry and Wildlife; and Smart Trees Pacific. We would also like to thank Dan Utter, members of Sönke Johnsen’s lab and attendees at Botany 2021 for useful feedback. This product uses the Census Bureau Data API but is not endorsed or certified by the Census Bureau.

## Author contributions

Conceptualization: DEM, BGS

Data Curation: DEM, BGS, WM, BFA, HK, MN, JK

Formal Analysis: DEM, BGS, WM

Funding acquisition: DEM, BGS, WM, BFA, HK, MN, JK

Investigation: DEM, BGS, WM, BFA, HK, MN, JK

Methodology: DEM, BGS, WM

Project administration: DEM, BGS

Resources: MN, JK

Software: DEM, BGS, WM

Supervision: DEM, BGS

Visualization: DEM

Writing – original draft: DEM, BGS

Writing – review & editing: DEM, BGS, WM, BFA, HK, MN, JK

## Data and materials availability

All data and code are available in the main text or the supplementary materials. The datasheets of city tree information from 63 cities (63 .csv files) have been uploaded to Dryad [after creating preprint with eLife, authors will insert the Dryad repository information].

## Supplementary Materials

Figs. S1 to S5

Tables S1 to S4

Data S1 to S9

## Supplementary Information

### This PDF file includes

Fig. S1 (supporting main-text Figure 2)

Fig. S2 (supporting main-text Figure 2)

Fig. S3 (supporting main-text Figure 2)

Fig. S4 (supporting main-text Figure 3)

Fig. S5 (supporting main-text Figure 4)

Table S1 (supporting main-text Figure 2)

Table S2 (supporting main-text Figure 4)

Table S3

Table S4

Legends for Datasets S1 to S9

### Other supplementary materials for this manuscript include the following

Datasets S1 to S9:

DataS1_City_Trees_Data_63_Files.zip

DataS2_Tree_Data_Summary_By_City.csv

DataS3_Calculate_Effective_Species.xlsx

DataS4_Code_for_Analysis_and_Plotting.zip

DataS5_Clustering_Results.csv

DataS6_Check_Native_Status_of_Species.xlsx

DataS7_Native_Taxa_By_State_BONAP.csv

DataS8_Environmental_PCA.xlsx

DataS9_Code_for_Data_Cleaning.zip

### Notes

- **Supporting figures**: in each main-text figure caption, I have indicated which supporting figures would be “child” figures to that main-text figure
- **Source Code and Data**: in each main text figure caption, I listed which source data and source code can be used to generate that figure

**Fig. S1.**
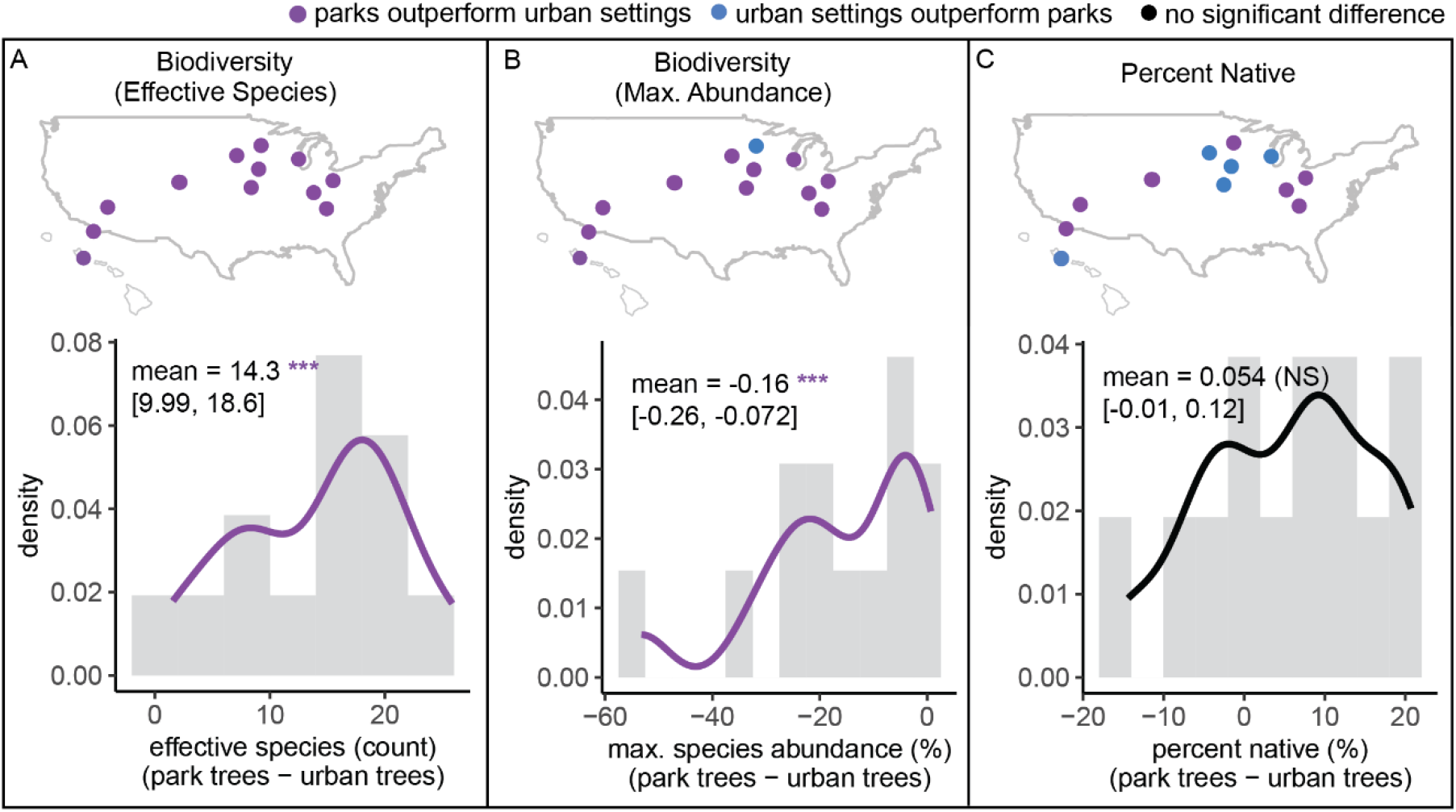
Tree communities in parks were significantly more biodiverse than those in urban settings but did not differ significantly in percent native. **(A)** Effective species numbers were greater in parks (two-sample paired t-test; t=7.2, p<0.0005, 95% CI=[9.99, 18.6], mean diff. = 14.3). **(B)** Maximum abundance of a single species was lower in parks (two-sample paired t-test; t=−3.9, p=0.0023, 95% CI=[-0.26, -0.072], mean diff. = −0.16). **(C)** Percent native did not differ significantly (two-sample paired t-test; t=1.8, p=0.092, 95% CI=[-0.01, 0.12], mean diff. = 0.054). We confirmed normality of the differences using a Shapiro-Wilk normality test, and confirmed all results through nonparametric Wilcoxon ranked-sign tests. We had sufficient data from 12 cities: Aurora, CO, park trees = 17366, urban trees N= 32455; Columbus, OH, park trees = 22791, urban trees N= 112586; Denver, CO, park trees = 252695, urban trees N= 12547; Des Moines, IA, park trees = 15035, urban trees N= 174; Honolulu, HI, park trees = 12650, urban trees N= 974; Knoxville, TN, park trees = 4821, urban trees N= 3551; Las Vegas, NV, park trees = 23636, urban trees N= 971; Louisville, KY, park trees = 16916, urban trees N= 4373; Milwaukee, WI, park trees = 18721, urban trees N= 28; Minneapolis, MN, park trees = 12983, urban trees N= 175232; Overland Park, KS, park trees = 162, urban trees N= 38898; San Diego, CA, park trees = 11260, urban trees N= 1012; and Sioux Falls, SD., park trees = 12672, urban trees N= 47684.

**Fig. S2.**
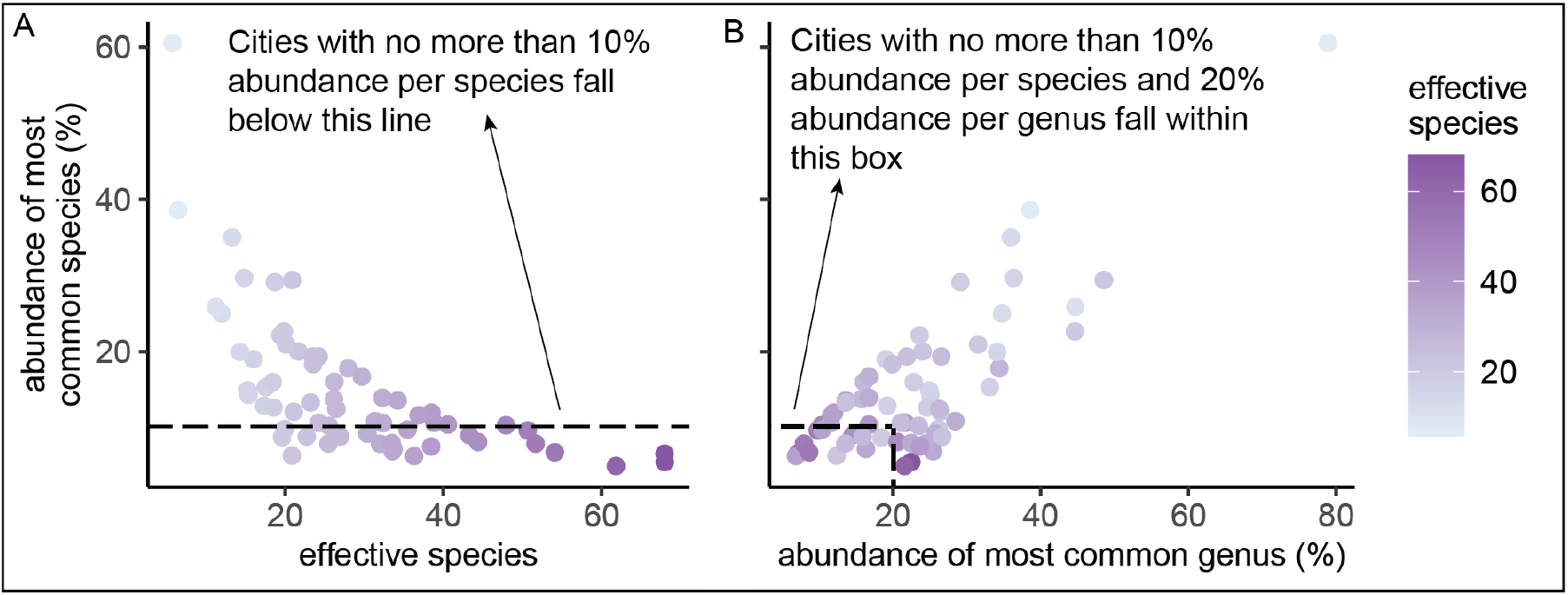
Effective species is a more nuanced metric of biodiversity than classic abundance-based measures. **(A)** The simple metric of abundance of most common species correlated with the more nuanced metric of effective species count (Pearson’s product-moment correlation: cor=−0.63, 95% CI=[-0.76, -0.46], t=−6.40, p<0.0005). **(B)** Abundance of the most common species versus abundance of the most common genus; most cities did not fulfill Santamour’s 10/20 rule.

**Fig. S3.**
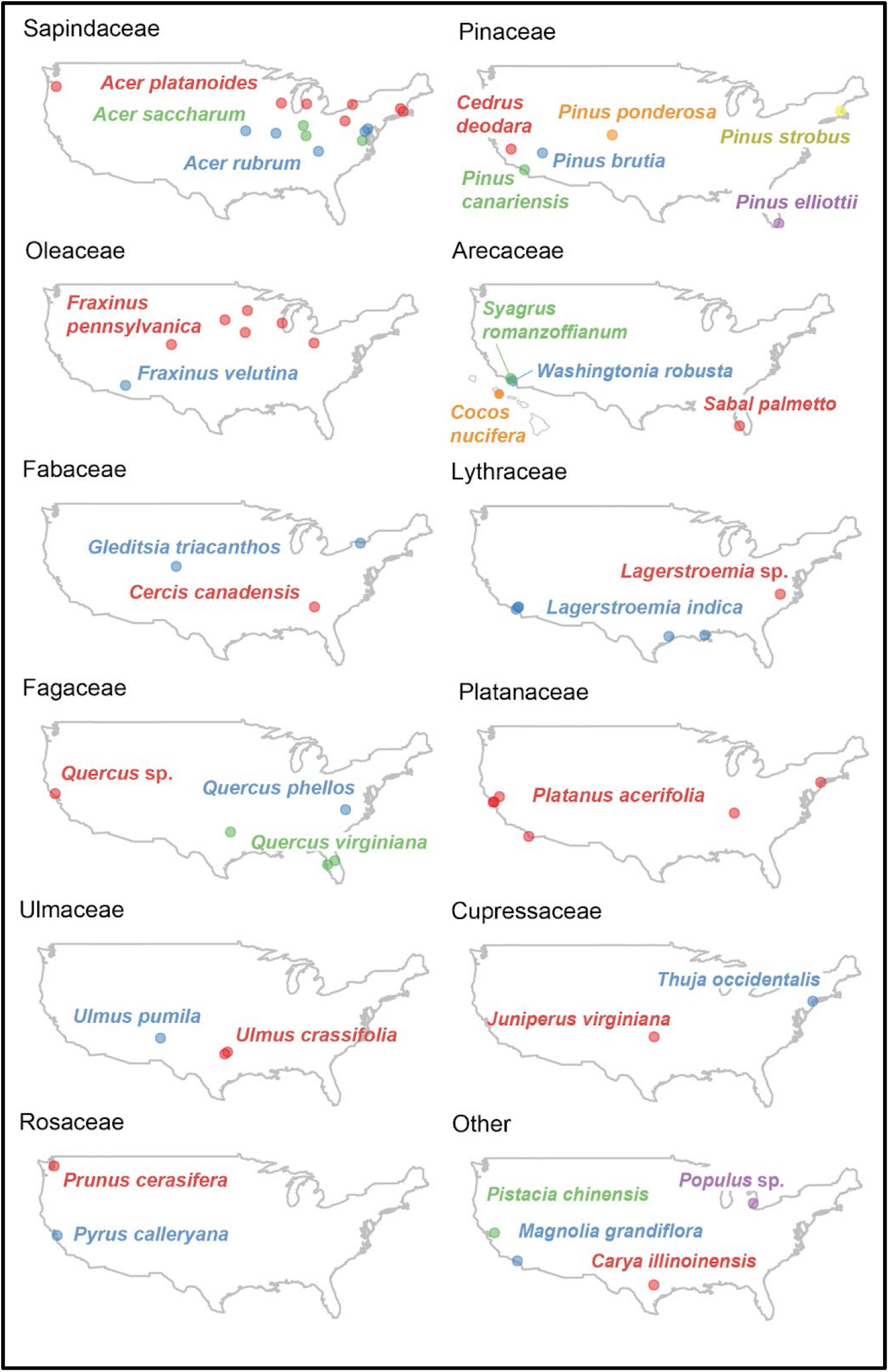
The most common species in each city is labeled (organized by family for display purposes).

**Fig. S4.**
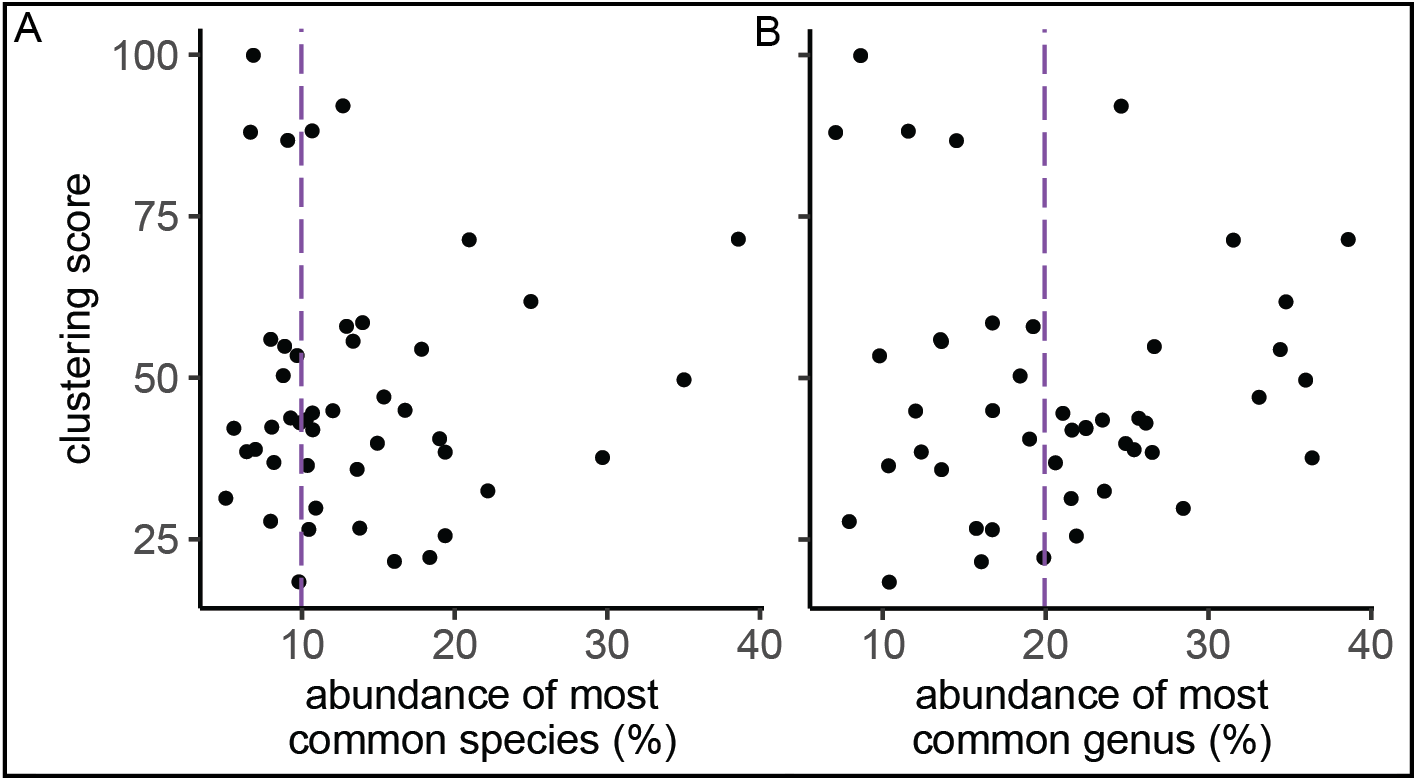
The degree to which a city has trees clustered by species does not correlate with abundance-based measures of biodiversity. Clustering score is the observed / expected effective species per cluster (see Methods and Main Text). **(A)** Clustering score versus abundance of the most common species. **(B)** Clustering score versus abundance of most common genus.

**Fig. S5.**
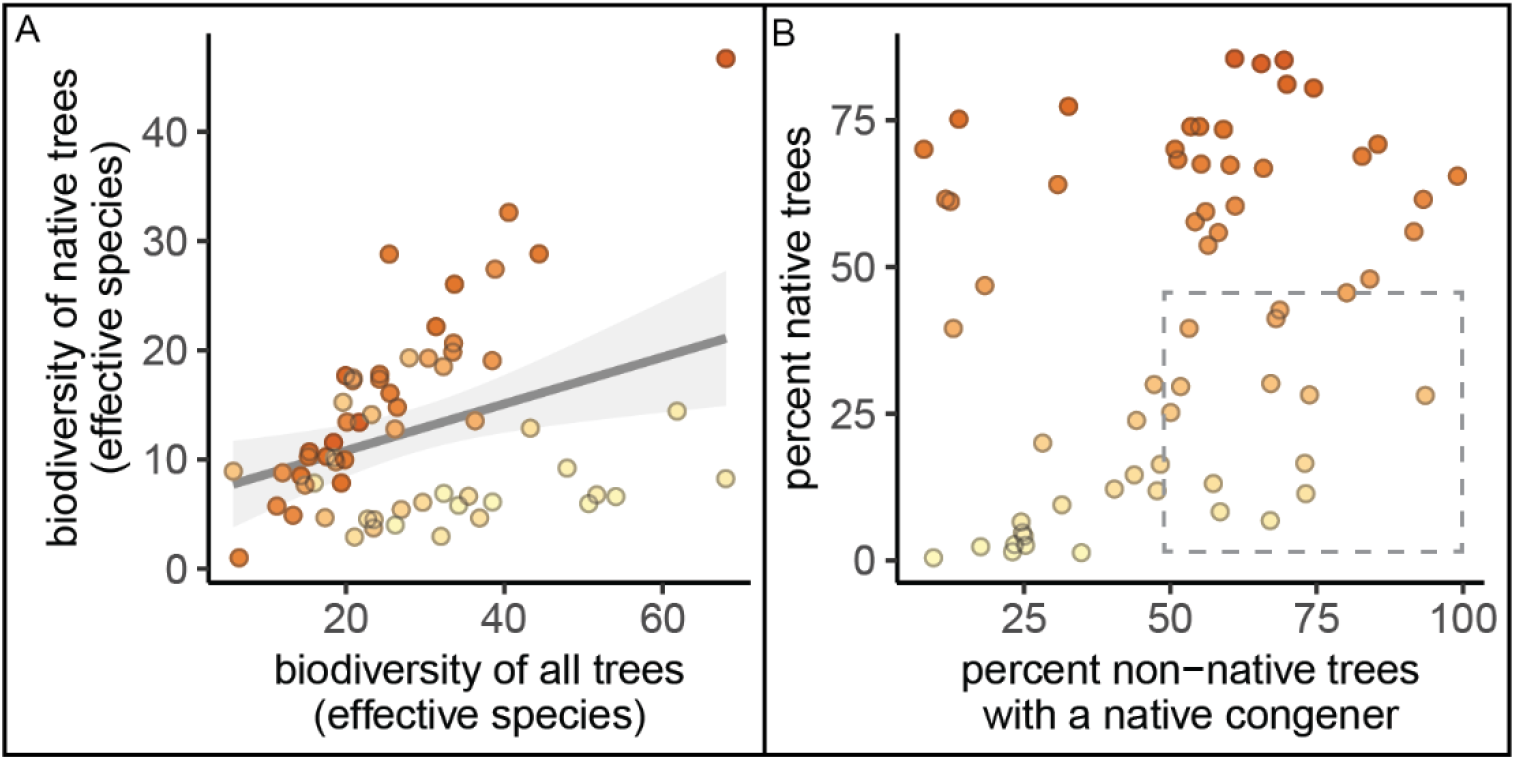
The percent of native trees is not the only important variable when considering nativity status; here we plot the biodiversity of native trees and percent of non-native trees that are closely related to native trees (have a native congener). **(A)** The biodiversity of all native trees (y-axis) is significantly correlated with biodiversity of all trees (Pearson’s product-moment correlation; cor = 0.35, 95% CI = [0.11, 0.55], p=0.0052); in both cases, we calculated effective species numbers. **(B)** The percent of all trees in a city that are native is not significantly related to the percent of all non-native trees that have a native congener. Cities with a low proportion of native trees (but high proportion of non-native trees with a native congener) are indicated with the grey dashed box.

**Table S1.** Here we report the results of linear regression models with effective species as the independent variable.

| Equation and Model Fit | Coefficient | All Data |  |  | Excluding outliers* |  |  |
| --- | --- | --- | --- | --- | --- | --- | --- |
|  |  | est. | 95% CI | p-val | est. | 95% CI | p-val |
| **effective_species ~**<br>**PC1:PC2 + PC1 +**<br>**log(number_trees)**<br>**All Data:**<br>AIC = 499.9 ; Adj. R<sup>2</sup>=0.20<br>**Excluding outliers:**<br>AIC = 479.4; Adj. R<sup>2</sup>=0.28 | PC1 | 1.65 | [0.49, 2.81] | 0.0061 | 0.91 | [-0.36, 2.18] | 0.16 |
|  | log(number_trees) | 6 | [1.64, 10.36] | 0.0078 | 8.19 | [3.65, 12.72] | 0.00064 |
|  | PC1:PC2 | -0.39 | [-1.01, 0.22] | 0.21 | -1.08 | [-1.89, -0.27] | 0.01 |
| **effective_species ~**<br>**PC1:PC2 +**<br>**log(number_trees)**<br>**All Data:**<br>AIC = 506.0; Adj. R<sup>2</sup> = 0.11<br>**Excluding Outliers:**<br>AIC = 479.6; Adj. R<sup>2</sup> = 0.26 | log(number_trees, 10) | 5.83 | [1.22, 10.44] | 0.014 | 8.6 | [4.06, 13.13] | 0.00036 |
|  | PC1:PC2 | -0.6 | [-1.24, 0.03] | 0.063 | -1.35 | [-2.07, -0.64] | 0.00037 |
| **effective_species ~**<br>**PC1:PC2 + PC1 +**<br>**tree_city_USA +**<br>**log(number_trees)**<br>**All Data:**<br>AIC = 500.0; Adj. R<sup>2</sup> = 0.21<br>**Excluding Outliers:**<br>AIC = 479.7; Adj. R<sup>2</sup> = 0.28 | PC1 | 1.89 | [0.68, 3.09] | 0.0027 | 1.14 | [-0.18, 2.45] | 0.09 |
|  | tree_city_USAyes | 7.33 | [-3.45, 18.11] | 0.18 | 6.57 | [-3.84, 16.97] | 0.21 |
|  | log(number_trees, 10) | 5.55 | [1.17, 9.93] | 0.014 | 7.72 | [3.15, 12.29] | 0.0013 |
|  | PC1:PC2 | -0.5 | [-1.13, 0.13] | 0.12 | -1.15 | [-1.97, -0.34] | 0.0064 |
| **effective_species ~**<br>**PC1:PC2 + PC1 + PC2 +**<br>**tree_city_USA +**<br>**log(number_trees)**<br>**All Data:**<br>AIC = 501.4; Adj. R<sup>2</sup> = 0.21<br>**Excluding Outliers:**<br>AIC = 481.67; Adj. R<sup>2</sup> = 0.27 | PC1 | 1.84 | [0.63, 3.06] | 0.0036 | 1.14 | [-0.19, 2.47] | 0.093 |
|  | PC2 | 0.49 | [-0.94, 1.91] | 0.5 | 0.12 | [-1.3, 1.53] | 0.87 |
|  | tree_city_USAyes | 6.94 | [-3.96, 17.83] | 0.21 | 6.48 | [-4.06, 17.03] | 0.22 |
|  | log(number_trees, 10) | 5.75 | [1.31, 10.18] | 0.012 | 7.73 | [3.12, 12.34] | 0.0015 |
|  | PC1:PC2 | -0.56 | [-1.23, 0.1] | 0.095 | -1.16 | [-1.99, -0.33] | 0.0068 |

\* Honolulu, HI and Miami, FL

**Table S2.**
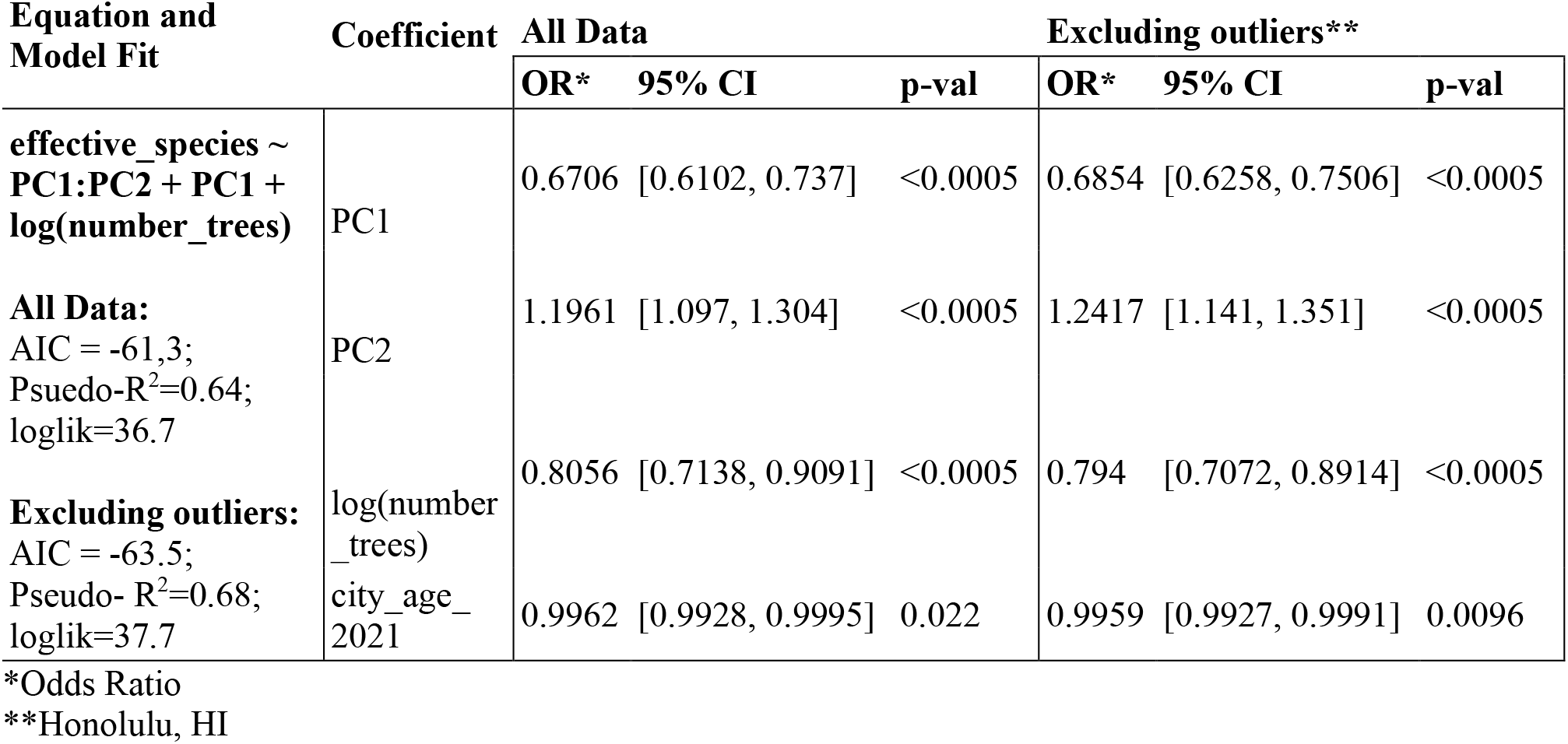
Here we report the results of beta regression models with percent native (percent of trees that are native) as the independent variable.

**Table S3.** We ran logistic regression models to identify correlations between condition and (i) tree size, (ii) tree location, and (iii) whether or not a tree was native. For size, smaller trees (lower diameter at breast height) tended to have better condition (9 of 12 cities). For location type, trees in the built environment tended to have better condition than those in parks (6 of 11 cities). For native status, results were mixed (native trees had no difference in condition for 8 of 15 cities, worse condition in 5 cities, and better condition in 2 cities).

|  | city | diameter at breast height (cm) |  |  | location type (park or urban setting) |  |  | native status |  |  | Model p- val |
| --- | --- | --- | --- | --- | --- | --- | --- | --- | --- | --- | --- |
|  |  | OR | sig | 95% CI | OR | sig | 95% CI | OR | sig | 95% CI |  |
| Native status, location type, diameter at breast height | Aurora, CO | 1.3 | * | [1.2,1.5] | 0.48 | * | [0.46,0.52] | 1.8 | * | [1.6,2] | <0.0005 |
|  | Columbus | 0.26 | * | [0.25,0.28] | 1.8 | * | [1.8,1.9] | 0.26 | * | [0.25,0.27] | <0.0005 |
|  | Knoxville | 0.2 | * | [0.15,0.27] | 0.66 | * | [0.56,0.77] | 0.61 | * | [0.51,0.73] | <0.0005 |
|  | SanDiego | 0.49 | * | [0.34,0.68] | 0.01 | * | [0.0064,0.016] | 0.09 | * | [0.071,0.11] | <0.0005 |
|  | Honolulu | 3.5 | * | [2,6.3] | 0.96 | NS | [0.57,1.5] | 0.59 | NS | [0.18,3.6] | <0.0005 |
|  | Louisville | 0.72 | * | [0.61,0.85] | 1.2 | * | [1.1,1.4] | 0.96 | NS | [0.83,1.1] | <0.0005 |
|  | Minneapolis | 48 | * | [40,57] | 1.2 | * | [1.1,1.4] | 0.99 | NS | [0.94,1] | <0.0005 |
|  | Sioux Falls | 0.049 | * | [0.035,0.068] | 0.12 | * | [0.11,0.13] | 0.93 | NS | [0.85,1] | <0.0005 |
| Native status, location type | Des Moines | NA | - | - | 0.82 | NS | [0.46,1.4] | 1.6 | * | [1.4,1.8] | <0.0005 |
|  | Las Vegas | NA | - | - | 0.1 | * | [0.0057,0.45] | 0.97 | NS | [0.87,1.1] | 0.0015 |
|  | Overland Park | NA | - | - | 0.048 | * | [0.019,0.16] | 1.1 | NS | [0.73,1.6] | 0.00025 |
| Native status, diameter at breast | Milwaukee | 0.57 | * | [0.44,0.74] | - | - | - | 0.59 | * | [0.51,0.69] | <0.0005 |
|  | Portland | 0.41 | * | [0.35,0.48] | - | - | - | 0.64 | * | [0.55,0.75] | <0.0005 |
|  | Richmond | 0.14 | * | [0.0358,0.548] | - | - | - | 1.2 | NS | [0.56,2.5] | 0.0084 |
|  | Seattle | 0.57 | * | [0.34,0.99] | - | - | - | 2.3 | NS | [0.95,7.8] | 0.032 |

**Table S4.**
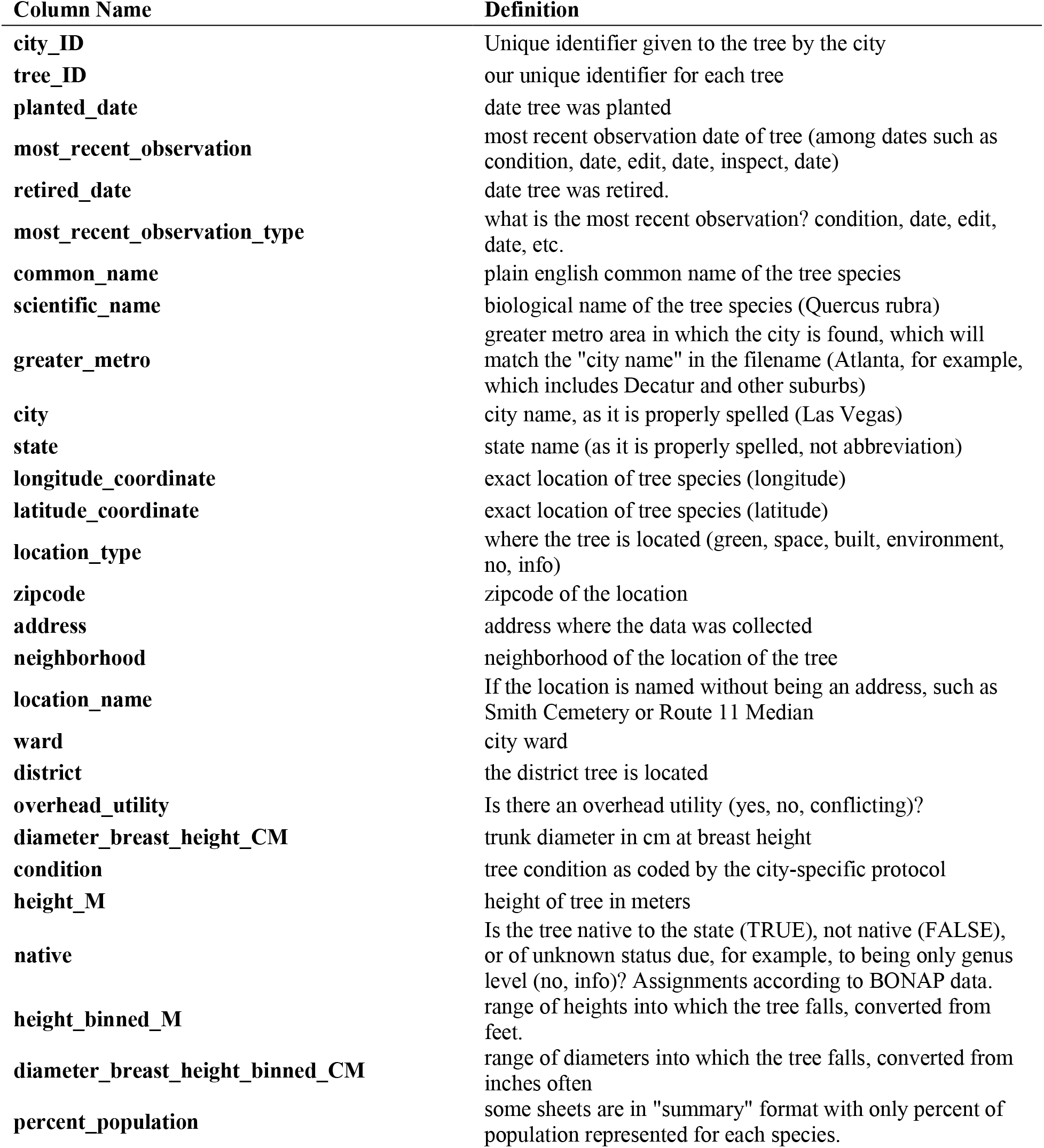
Here we define the standardized columns used herein.

| Column Name | Definition |
| --- | --- |
| **city_ID** | Unique identifier given to the tree by the city |
| **tree_ID** | our unique identifier for each tree |
| **planted_date** | date tree was planted |
| **most_recent_observation** | most recent observation date of tree (among dates such as condition, date, edit, date, inspect, date) |
| **retired_date** | date tree was retired. |
| **most_recent_observation_type** | what is the most recent observation? condition, date, edit, date, etc. |
| **common_name** | plain english common name of the tree species |
| **scientific_name** | biological name of the tree species (Quercus rubra) |
| **greater_metro** | greater metro area in which the city is found, which will match the "city name" in the filename (Atlanta, for example, which includes Decatur and other suburbs) |
| **city** | city name, as it is properly spelled (Las Vegas) |
| **state** | state name (as it is properly spelled, not abbreviation) |
| **longitude_coordinate** | exact location of tree species (longitude) |
| **latitude_coordinate** | exact location of tree species (latitude) |
| **location_type** | where the tree is located (green, space, built, environment, no, info) |
| **zipcode** | zipcode of the location |
| **address** | address where the data was collected |
| **neighborhood** | neighborhood of the location of the tree |
| **location_name** | If the location is named without being an address, such as Smith Cemetery or Route 11 Median |
| **ward** | city ward |
| **district** | the district tree is located |
| **overhead_utility** | Is there an overhead utility (yes, no, conflicting)? |
| **diameter_breast_height_CM** | trunk diameter in cm at breast height |
| **condition** | tree condition as coded by the city-specific protocol |
| **height_M** | height of tree in meters |
| **native** | Is the tree native to the state (TRUE), not native (FALSE), or of unknown status due, for example, to being only genus level (no, info)? Assignments according to BONAP data. |
| **height_binned_M** | range of heights into which the tree falls, converted from feet. |
| **diameter_breast_height_binned_CM** | range of diameters into which the tree falls, converted from inches often |
| **percent_population** | some sheets are in "summary" format with only percent of population represented for each species. |

**Data S1. (separate file; Data_S1_City_Trees_Data_63_Files)**. This zipped file includes all of the cleaned data used in the study (63 spreadsheets, one for each city, where each row is a tree).

**Data S2. (separate file; DataS2_Tree_Data_Summary_By_City.csv)**. Here we present all results by city, including number of trees, percent native, effective species count, environmental variables, socio-cultural variables, and more.

**Data S3. (separate file; DataS3_Calculate_Effective_Species.xlsx)**. We developed an Excel Spreadsheet which calculates effective species counts, a robust measure of biodiversity, from (i) a list of all trees or (ii) a list of species counts.

**Data S4. (separate file; DataS4_Code_for_Analysis_and_Plotting)**. This zipped file includes all code used to analyze and plot the results reported in this paper.

**Data S5. (separate file; DataS5_Clustering_Results.csv)**. This file includes all statistics on clustering by species, including number of clusters, median effective species count per cluster, min, max, and inter-quartile range (see Fig. 2).

**Data S6. (separate file, DataS6_Check_Native_Status_of_Species.xlsx)**. This Excel Workbook allows readers to input their list of species, select a state, and receive a corresponding list of whether or not each species is native to that state (based on BONAP designations).

**Data S7. (separate file, DataS7_Native_Taxa_By_State_BONAP.csv)**. This csv file includes a list of all native taxa observed in each US state, from the Biota of North America Project.

**Data S8. (separate file; DataS8_Environmental_PCA.xlsx)**. This file includes all loadings and scores for the environmental PCA (see Fig. 3A).

**Data S9. (separate file; DataS9_Code_for_Data_Cleaning)**. This zipped file includes all code sheets in Python and R, and instructions, for the full data cleaning procedure (See Methods, Data Cleaning).

